# Macropinocytosis of aggregated amyloid-beta and tau requires Arf6 and the RhoGTPases Rac1, Cdc42 and RhoA

**DOI:** 10.1101/2025.03.02.641073

**Authors:** Jordan M. Krupa, Manoj Reddy Medapati, Adrianna Tsang, Claudia Seah, Stephen H. Pasternak

## Abstract

The neuron-to-neuron transfer of amyloid-beta (Aβ) and tau aggregates have been proposed to underlie the propagation of protein aggregation in Alzheimers disease (AD) and contributing to progressive neurodegeneration. Several studies have provided evidence that aggregates of Aβ and tau are taken up into neuronal cells and neurons from the extracellular environment, where they contribute to the propagation of Aβ and tau aggregation in AD through seeding their aggregation. As a result, attention has been placed on determining the cellular mechanisms that contribute to this uptake, with the hopes that targeting these mechanisms could halt the progression of AD by preventing aggregate transfer. Previous studies have demonstrated the uptake of Aβ and tau aggregates through the endocytic process called macropinocytosis. The activity of several GTPases has been demonstrated to regulate macropinocytosis, including Arf6 and the RhoGTPases Rac1, Cdc42 and RhoA. Here, we examined the uptake of Aβ42 oligomers and tau fibrils by macropinocytosis in neurons and the role of Arf6, Rac1, Cdc42 and RhoA activity in the macropinocytosis of these aggregates. In this study, we demonstrated that extracellular Aβ42 oligomers and tau fibrils are taken up by iPSC-derived neurons and delivered directly to LAMP1- labeled lysosomes through macropinocytosis. This was demonstrated by reduced uptake in response to treatment with the macropinocytosis inhibitor EIPA, but not in response to the clathrin inhibitor Pitstop2. The uptake of these aggregates by macropinocytosis was significantly reduced by the inhibition of the GTPases Arf6, Rac1, Cdc42 and RhoA. Further, we also demonstrated that accumulation of extracellular Aβ42 oligomers and tau fibrils results in an accumulation of cytoplasmic tau within aggregate-containing lysosomes. Together these results provide evidence that the GTPases Arf6, Rac1, Cdc42 and RhoA play a role in the neuronal uptake of Abeta and tau aggregates by macropinocytosis and identifies new molecular targets to explore preventing the neuron-to-neuron transfer of these aggregates in AD.

## Introduction

Progressive neurodegenerative disorders affect many Canadians and pose a significant healthcare and economic challenge. These disorders are characterized by the dysfunction and degeneration of neurons within the central nervous system. The degeneration of neurons can be tracked into discrete stages, which parallel the progressive loss of cognitive functions (1). The most common of these disorders, Alzheimer’s disease (AD), affects a significant portion of the population and poses a critical challenge for our healthcare system as our population ages (2). Thus, the urgent need to develop new therapies to combat this disease is abundantly clear.

AD diagnosis begins in a prodromal stage known as mild cognitive impairment (MCI), where subtle changes in cognition can be noted (3). This begins with loss of neurons within the medial temporal lobe, specifically within the entorhinal cortex and hippocampus, and then spreading throughout the neocortex (1). These individuals develop significant impairments to their memory and attention, and then develop difficulties with language, frontal executive function and visuospatial function (4). The disease eventually progresses to the point where individuals cannot perform daily life functions and require extensive care.

Underlying this dysfunction and loss of neurons is the accumulation of toxic aggregates in the brain. These include the small peptide amyloid-beta (Aβ) and the microtubule-binding protein tau. Our current understanding is that Aβ produced by the enzymatic cleavage of amyloid precursor protein (APP) begins to aggregate in the brains of patients long before the development of symptoms (5–7). Continued aggregation of Aβ is thought to disrupt synapses within specific brain regions, contributing to and triggering the seeding of tau aggregation (8–10). Tau aggregation is thought to contribute further to the seeding of tau aggregates, neuronal dysfunction and causes neuronal death (11). These events are thought to first occur in discrete vulnerable brain areas such as the hippocampus, then spread through interconnected regions (12). This phenomenon is thought to be perpetuated by the transcellular or neuron-to-neuron transfer of aggregates between either connected or neighboring neurons (13,14).

The transcellular spread of toxic protein aggregates requires two things: the release of the toxic aggregate and the uptake of that aggregate into a naïve cell (14). Preventing the release of toxic aggregates from neurons could be explored to prevent transcellular spread. However, because these neurons have already developed pathology and ultimately will undergo cell death, the release of aggregates may be inevitable regardless of intervention (14). Given this, preventing the uptake of toxic aggregates into healthy neurons might prevent the seeding of aggregation and be a therapeutic target.

Uptake of aggregates requires the endocytosis of the proteins from the extracellular space into the cell. A cell may use several different endocytic mechanisms to take up external materials. Macropinocytosis, or ‘cellular drinking,’ is an endocytic mechanism in which a cell extends its plasma membrane to engulf a portion of extracellular fluid and material (15). These extensions form circular cups and then fuse and detach from the plasma membrane, creating macropinosomes (16–18). These then quickly fuse with lysosomes, resulting in rapid internalization of extracellular material (19). This process stands in contrast with more common endocytic pathways like clathrin-mediated endocytosis (CME) (20). Macropinocytosis occurs much more rapidly, can internalize larger particles, and results in immediate delivery to lysosomes (21,22). Given the large size of Aβ and tau aggregates and their presence in extracellular fluid in AD patients, macropinocytosis appears to be a good candidate for the uptake of aggregates (23–25).

A handful of studies have previously investigated the internalization of Aβ and tau. Aβ oligomers have been observed to internalize rapidly into cells when added to cultures of immortal cell lines (26). In addition, they observed that this process was both heparan sulfate proteoglycan (HSPG) and cholesterol-dependent (26), characteristic of macropinocytosis. Additionally, inhibition of CME resulted in no reduction of Aβ uptake (26). Similarly, tau pre-formed fibril (PFF) uptake was also observed to be HSPG dependent and occurred rapidly (27). While there are now a handful of studies demonstrating macropinocytosis of Aβ and tau aggregates, there are currently no studies that directly link the uptake of Aβ and tau aggregates to known macropinocytic regulators. Several small GTPases known as Arf6, Rac1, RhoA and Cdc42, key proteins in the modulation of actin dynamics, have been demonstrated to regulate macropinocytosis (28). Given these observations, the objective of the current study was to examine the role of macropinocytosis in Aβ and tau aggregate uptake, examine the roles of GTPases in its regulation, and elucidate the contributions of the macropinocytosis of aggregates to the seeding of intracellular tau aggregation. Here we demonstrated that both Aβ42 oligomers (Aβ42o) and tau fibrils are taken up into human neurons derived from iPSCs by macropinocytosis and delivered to lysosomes. This uptake was reduced following treatment with EIPA or inhibition of Arf6, Rac1, Cdc42 and RhoA. Additionally, incubating cells with Aβ42o or tau fibrils resulted in an increase in the amount of active GTP-bound Rac1, Cdc42 and RhoA. The uptake of these aggregates by macropinocytosis and their delivery to lysosomes resulted in an accumulation of intracellular tau into swollen lysosomes containing exogenous Aβ42o and/or tau fibrils.

## Methods

### Cell culture materials and methods

The Arf6 inhibitor NAV-2729 (SML2238), Rac1 inhibitor EHT 1864 (E1657), and Cdc42 inhibitor (SML0407), 1,1,1,3,3,3-Hexafluoroisopropanol (HFIP) were purchased from Sigma-Aldrich (MO, USA). The RhoA selective inhibitor Rhosin hydrochloride (5003) and Ethylisopropyl amiloride (EIPA; 3378) were purchased from Tocris (5003; Bristol, UK). Pitstop2 was purchased from abcam (ab120687; MA, USA). G-LISA kits for GTP-bound Rac1 (BK128), Cdc42 (BK127) and RhoA (BK124) were all purchased from Cytoskeleton (CO, USA). Halo-tag ligand labeled with Janelia Fluor (JF) 549 (GA1110) and 646 (GA1120) were obtained from Promega (WI, USA). Lysine fixable 70 kDa dextran labeled with tetramethylrhodamine (TMR) was purchased from Invitrogen (D1818; MA, USA)

Human induced pluripotent stem cells (iPSCs) were obtained from the New York Stem Cell Foundation (NYSCF; BN0004-01-CS-001; Lot# 241-1D; Parent cell line ID: 7889B/7889O; sex: male). Cell culture reagents: Dulbecco’s phosphate-buffered saline (DPBS, 14190144), Hank’s balanced salt solution (HBSS, 14025092) was purchased from Gibco (CA, USA); dimethyl sulfoxide (DMSO, D8418) from Sigma-Aldrich (MO, USA). 50X B27 supplement (17504044), L-glutamine supplement alternative GlutaMAX (35050–061), human recombinant BDNF (450–02), human recombinant GDNF (450–10), and antibiotic-antimyotic (15240096) were all purchased from Gibco (CA, USA). Dibutyryl cyclic-AMP (db-cAMP) was purchased from Sigma-Aldrich (MO, USA).

### Viral vectors and DNA constructs

The following constructs were produced and packaged into adenovirus serotype 5 (Ad5) replication incompetent viruses by VectorBuilder (IL, USA); LAMP1-mCherry (LAMP1-mCh), LAMP1-TagBFP2 (LAMP1-BFP), Rab5-mCherry (Rab5-mCh), PLCδPH-TagBFP2 (PLCδPH-BFP), and Halo-tag fused tau containing a P301L mutation (Halo-tau). P301L mutant tau was used as it shows an increased propensity to aggregate due to reduced microtubule binding compared to wild type (29). IPSC-derived neurons were transduced with each of these viruses at an MOI of 50.

### Neural progenitor cell culture and IPSC-derived neuron differentiation, maturation, and infection

The induction and expansion of human iPSCs (NYSCF) to CNS-type neural progenitor cells (NPCs) was performed by our laboratory technician Clauda Seah using StemCELL Technologies STEMdiff SMADi Neural Induction Kit (08581), using their STEMdiff neural induction media (05835), following manufacturer protocols (Technical Manual; Version 03; Document # 10000005588; STEMCELL technologies; CA). For maintenance of cultured NPCs, cells were cultured and passaged every 5-7 days in STEMdiff neural progenitor medium (05834; STEMCELL technologies; CA) in 6-well plates coated with poly-L-ornithine (PLO; 15 ug/mL) and laminin (10 ug/mL).

For the differentiation of NPCs to IPSC-derived neurons, NPCs were plated at the time of passaging and cultured in STEMdiff Forebrain Neuron Differentiation media (08601; STEMCELL technologies; CA) on PLO and laminin-coated culture ware for 7 days with media changed every 24 hours. After differentiation, neurons were plated (day in-vitro; DIV0) at a density of 1.50 x 10^4^ cells on PLO- and laminin-coated 35mm dish with a 14mm glass-bottom (P35G-1.5-14-C, MatTek, MA, USA). Maturation of differentiated neurons occurred up to DIV21 in neurobasal medium (21103049; Gibco; CA, USA) supplemented with B27 (1X), GlutaMAX (0.5X), db- cAMP (0.5 mM), BDNF (20 ng/mL), GDNF (20 ng/mL), ascorbic acid (0.2 ug/mL) and antibiotic-antimycotic (1X) (protocol adapted from Stem Cell Technologies). During maturation, 50% of the media on dishes was replaced with fresh-supplemented media. Due to the experiments in this study being conducted in live cells, successful generation of IPSC-derived neurons was assessed in separate dishes plated for differentiation quantification alongside cells plated for experiments. To assess the differentiation of NPCs to IPSC-derived neurons, plates were stained for the neuron marker MAP2 and the NPC marker Nestin along with nuclear staining by DAPI. The number of MAP2+ and Nestin+ cells were assessed to ensure the majority of cells differentiated to IPSC-derived neurons. For selection of neurons used in live cell imaging experiments, cells were selected based upon morphology which matched that of MAP2 stained cells. Infections of cells were performed at DIV18, using the number of cells plated at DIV0 to calculate the desired MOI of a given viral vector. Viral vector was added directly to 1 mL of existing media and was incubated with cells for 48 hours.

### Aβ42 oligomer and tau pre-formed fibril preparation

Recombinant Aβ42 tagged with HyLyte Fluor 647 (Anaspec; AS-64161; CA, USA) was used to produce Aβ42 oligomers (Aβ42o) following previously established protocols (30,31). Briefly, lyophilized Aβ peptides were first solubilized in HFIP to a concentration of 1 mM. HFIP was removed by vacuum centrifugation at room temperature, yielding a dry peptide film. To produce Aβ42o, the peptide film was dissolved in ice-cold phenol-free F-12 media to a concentration of 100 μM and incubated for 24 hrs at 4°C. PFFs were generated from recombinant human tau (2N4R) with a P301S mutation tagged with ATTO 488 (StressMarq; SPR-329B-A488; Canada) following manufacturers sonication protocol.

### Inhibitor treatments and aggregate internalization experiments

EIPA is a commonly used inhibitor of macropinocytosis, through its effect on Rac1 and Cdc42 activity due to plasma membrane pH changes produced by blocking the activity of Na+/H+ exchangers (32). Pitstop2 is an inhibitor of clathrin-coated pit formation (33) and was used to inhibit CME. This was chosen for this study over dynamin inhibition using Dynasore because it has been shown to have off-target effects resulting in the inhibition of dynamin-independent endocytosis, including macropinocytosis (34). To assess the contributions of macropinocytosis and CME to Aβ42o and tauPFF internalization to lysosomes, LAMP1-mCh infected DIV21 neurons were treated with 0.1% DMSO for 30 minutes, 10 μM EIPA for 30 minutes, or 10 μM Pitstop2 for 5 minutes prior to incubation with either aggregate. Following these treatments, cells were incubated with 500 nM of Aβ42o or 5 μg/mL of tauPFFs in HBSS for 30 mins, 1 hr, or 2 hrs. After incubation, plates were washed with warm HBSS to remove aggregates not taken up and immediately moved to the microscope for live cell imaging. This resulted in the remaining signal in the dish from either Aβ42o or tauPFFs being either aggregates taken up into cells or aggregates stuck to the coating on the glass dish.

To assess the roles of small GTPases in Aβ42o and tauPFF uptake, the inhibitors NAV- 2729 (Arf6), EHT 1864 (Rac1), ML 141 (Cdc42), and Rhosin (RhoA) were used. As a vehicle control, 0.1% DMSO (v/v) was used. Prior to incubation of DIV21 neurons with aggregates, cells were treated at the following concentrations: 5 μM NAV-2729 (3 hrs), 20 μM EHT 1864 (18 hrs), 20 μM ML 141, 35 μM Rhosin (3 hrs), and 0.1% DMSO (18 hrs). EHT 1864 and ML 141 concentrations were used in our previous study (35), and NAV-2729 and Rhosin treatments were based on previous literature using these inhibitors (36,37). Cells were infected with either LAMP1- mCh or LAMP1-BFP and treated as described above were then incubated with 500 nM of Aβ42o or 5 μg/mL of tauPFFs in HBSS for 30 mins, then washed with warm HBSS and immediately imaged live. For experiments examining co-uptake with Dex70, 1 mg/mL of dextran 70kDa was added along with Aβ42o or tauPFF.

For qualitative experiments assessing the presence of Aβ42o or tauPFFs in membrane ruffles, neurons transduced with PLCδPH-BFP were incubated with either aggregate at the above concentrations for 10 minutes then subsequently fixed in 4% paraformaldehyde (PFA) for 10 mins at room temperature. After fixation F-actin was labelled using the fluorescently tagged phalloidin called Acti-Stain 555 (Cytoskeleton; PHDH1-A; CO, USA) according to manufacturer protocol.

### Halo-tau pulse chase experiment

To assess the effects of Aβ42o and tauPFF uptake on the localization of cytoplasmic tau, neurons were infected with Halo-tau and LAMP1-BFP. After infection (DIV20), cells were incubated overnight with no aggregates, 100 nM of Aβ42o, 1 μg/mL of tauPFFs, or both Aβ42o and tauPFFs together. Following this, Halo-tau was labelled using JF549 (Aβ42o alone and both aggregates together) or JF646 (untreated and tauPFFs) labelled Halo-tag ligands according to manufacturer protocols. After labelling Halo-tau, neurons were imaged live directly following labelling (0 hrs), or at 6 hrs, 24 hrs, 48 hrs, 5 days, and 7 days after labeling.

### Confocal microscopy

Live cell imaging was performed using a Leica DMI8 STELLARIS inverted confocal microscope, using an HC PL APO CS2 63X 1.4 numerical aperture oil immersion lens. The optical section thickness was typically 1 micron. TagBFP2 was imaged using a 405 nm laser with a 415- 560 nm filter set; ATTO 488 fluorescence was imaged using a 488 nm laser line with a 500-550 nm filter set on a HyD hybrid detector; JF549 and mCherry were imaged using a 542 nm laser line with a 570-620 nm filter set; JF656 and HyLyte 647 were imaged using a 638 nm laser line with a 650-700 nm filter set. Images of the cell body of individual neurons were taken as z-stacks to capture the entirety of cells.

### Data quantification and analysis

Analysis of all images was performed using Imaris 10.1.1 imaging software (Bitplane, CT, USA). For experiments examining the uptake of Aβ42o and tauPFFs to lysosomes and early endosomes, colocalization between Aβ42o or Tau with LAMP1 and Rab5 was measured. We used a similar technique to what our lab has previously established (38), setting the colocalization threshold of Aβ42o-647 or tauPFF-488 signal to include the brightest 0.5% of the total signal in every image analyzed. Thresholding for LAMP1 and Rab5 was determined using the brightest signal which represented a clear punctate compartment. In doing so, we measured the percentage of Aβ42o and tauPFF signal above the set threshold colocalized with LAMP1 or Rab5 in an unbiased manner, allowing for the assessment of changes in the amount of aggregate in these compartments over time or between different treatments. This same approach was used in the measurement of colocalization between Aβ42o or tauPFFs with LAMP1 in cells treated with inhibitors. To assess the co-uptake of Aβ42o and tauPFFs with Dex70, the spots function in Imaris was used to mark Aβ42o or tauPPF puncta and Dex70 puncta. Doing so, the number of puncta contained within each cell body was quantified, along with the number of puncta containing aggregates and Dex70 together. We then determined the number of these puncta in neurons treated with vehicle control. Only spots ranging from 0.2-0.6 μm were considered for analysis as this the typical size of a macropinosomes. Graphing and statistical analysis were performed using GraphPad Prism 10. Statistical tests varied depending on the experiment, but for all tests, p-values of less than 0.05 were considered statistically significant.

### Neuro2a cell culture and Rac1, Cdc42 and RhoA G-LISA assays

Neuro-2a (N2a) mouse neuroblastoma cells were acquired from ATTCC (CCL-131, VA, USA) and were utilized for G-LISA assays rather than IPSC-derived neurons to obtain enough protein to run the assay within the linear range. N2a cells were cultured in MEM containing 10% FBS in a 25 cm^2^ flask (130189, Thermo Scientific, MA, USA) at 5% CO2 at 37 °C. Cells were passaged by trypsinization every 3-4 days. One day before transfection, cells were seeded at a density of 5.0 x 10^4^ cells in each well of a 4-well plate. Cells were subsequently transfected with the APP695 construct using lipofectamine 2000 (11668019, Invitrogen, MA, USA) according to manufacturer protocol. The following day, the cells were serum starved overnight for 18 hours to ensure low basal levels of GTPase activity and induce differentiation. Following serum starvation, cells were placed on ice, and 4-well plates were treated with 500 nM Aβ42o, 5 μg/mL tauPFFs or HBSS (no-treatment control). As a positive control, a 4-well plate was treated with EGF. Each 4- well plate was incubated with its respective treatment for 20 minutes on ice and then incubated at 5% CO2, 37 °C for 5 minutes. Plates were immediately moved back to ice, washed two times with ice-cold PBS, and then protein was extracted according to manufacturer protocol. Protein concentration was measured by BCA to ensure equivalent amounts of total protein were loaded, and G-LISAs were then run according to manufacturer protocol in technical triplicates. Reagents used for N2a culture include: minimum essential media (MEM, 11095080), Dulbecco’s phosphate-buffered saline (DPBS, 14190144), Hank’s balanced salt solution (HBSS, 14025092) were purchased from Gibco (CA, USA); fetal bovine serum (FBS, 090-150) from Wisent (QC, Canada).

## Results

### Amyloid-beta oligomers and tau fibrils are rapidly internalized to lysosomes

To begin investigating the macropinocytosis of Aβ and tau aggregates, we first characterized the trafficking of Aβ42o and tau-PFFs, to early endosomes and lysosomes. Delivery rapidly to lysosomes is a commonly observed endpoint for macropinocytic cargo, whereas cargo internalized by CME is typically trafficked to early endosomes followed by late endosomes before cargo is delivered to lysosomes. To investigate the uptake of oligomers and fibrils to early endosomes and lysosomes, cultured human neurons were transduced with constructs containing LAMP1-BFP to label lysosomes and Rab5-mCh to label early endosomes. Cells were then incubated with fluorescently tagged Aβ42o or tau-PFFs for 15 mins, 30 mins, 1 hr, and 2 hrs. Cells were then immediately imaged live, and colocalization with both compartment markers was quantitated using Imaris 10.1 software (Bitplane). Pixels containing colocalized markers are indicated in white. Aβ42o was observed in both Rab5 and LAMP1 labeled compartments, with more colocalization being seen with LAMP1 (Figure 1A). While colocalization increased with LAMP1 over time, no changes were observed with Rab5 (Figure 1A). Colocalization of Aβ42o with LAMP1 was significantly greater than Rab5 at 1 hr (6.981 ± 1.568; p=0.008) and 2 hrs (11.58 ± 1.568; p<0.001; Figure 1B). TauPFFs also showed more colocalization with LAMP1, which increased with longer incubation times (Figure 1C). Significantly higher colocalization with LAMP1 was observed at 30 min (13.17 ± 4.009; p=0.03), 1 hr (19.47 ± 4.91; p=0.007) and 2 hrs (21.53 ± 4.009; p<0.001; Figure 1D).

**Figure 1.**
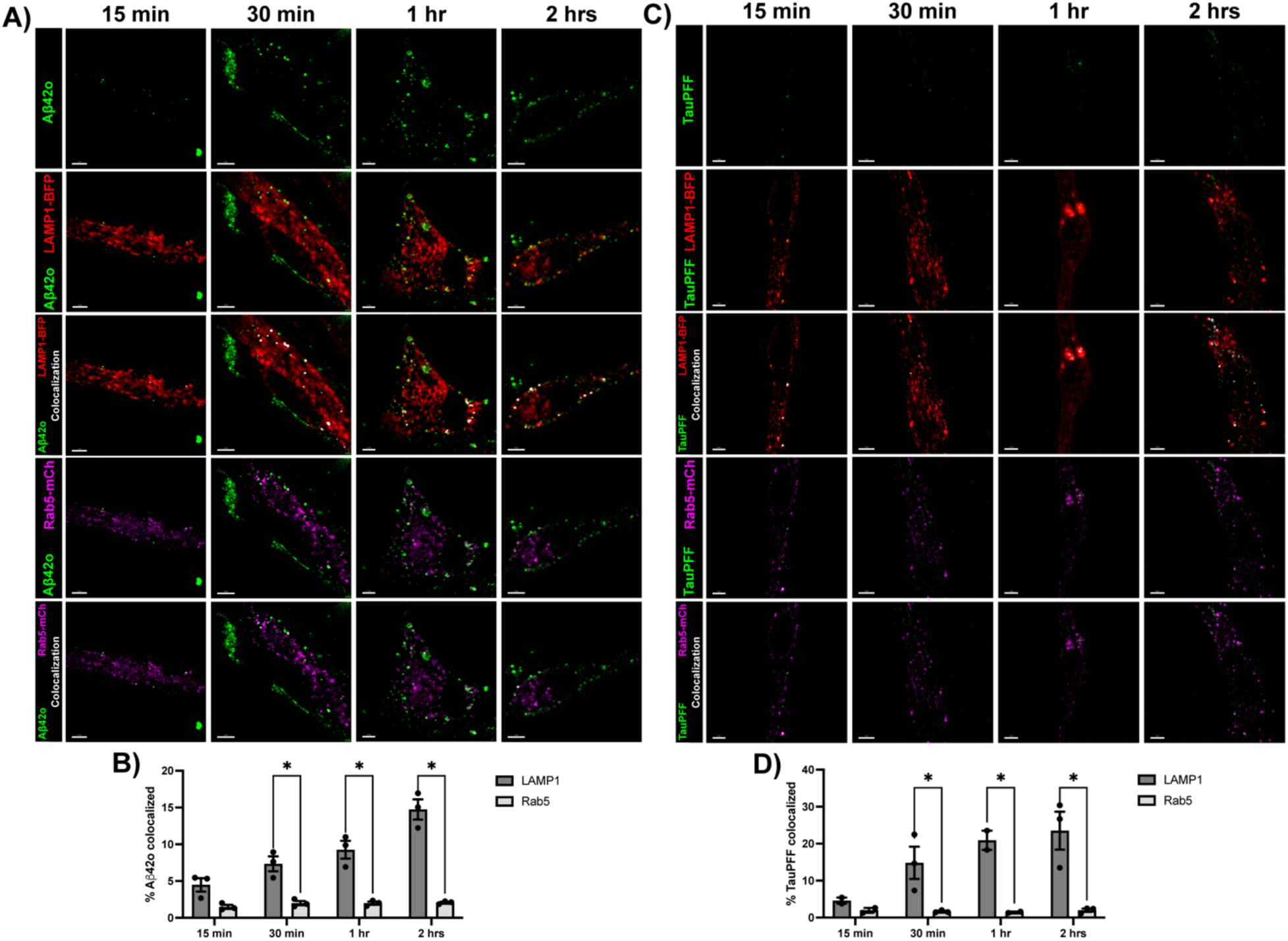
AB42o and tauPFFs are rapidly trafficked to lysosomes. **A)** IPSC-derived neurons were transduced with LAMP1-BFP (red) and Rab5-mCh (magenta) and were incubated with Aβ42o (red) for 15 min, 30 min, 1 hr, and 2 hrs. Colocalization between Aβ42o and each marker was quantitated using Imaris 10.1 software and is denoted by white pixels. **B)** Quantification of mean percent of Aβ42o signal colocalized with LAMP1 or Rab5 from three replicate experiments (n=3), each containing at least 6 individual neuron cell body images at each time point. Difference between LAMP1 and Rab5 was assessed using a two-way ANOVA with Sidak’s test. **C)** Transduced neurons were incubated with tauPFFs (green) for 15 min, 30 min, 1 hr, and 2 hrs. Colocalization between tauPFF and LAMP1 or Rab5 is marked by white pixels. **D)** Quantification of the mean percent of tauPFF colocalized with LAMP1 or Rab5 from three replicate experiments (n=3), each containing at least 6 individual neuron cell body images at each time point. Difference between LAMP1 and Rab5 was assessed using a two-way ANOVA with Sidak’s test. *Data is presented as mean* ± *SEM. * p<0.05. Scale bar = 5μm*.

### Inhibition of macropinocytosis prevents the rapid uptake of aggregates to lysosomes

We next examined the effect of the macropinocytosis inhibitor EIPA and the CME inhibitor Pitstop2 on the uptake of Aβ42o and tauPFFs. Prior to incubation with either aggregate, LAMP1- BFP expressing treated neurons were then incubated with fluorescent-tagged Aβ42o or tauPFFs for 30 mins, 1 hr, or 2 hrs, then imaged live and colocalization with LAMP1-labeled lysosomes was quantitated. Internalization of Aβ42o to LAMP1-labeled compartments was observed with either vehicle control or Pitstop2 treatment, but colocalization was dramatically inhibited by EIPA at all time points (Figure 2A). The percentage of Aβ42o colocalized with LAMP1 was significantly reduced with EIPA treatment compared to DMSO at 30 mins (-4.776 ± 1.078; p<0.001), 1 hr (- 7.427 ± 1.078; p<0.001) and 2 hrs (-12.27 ± 1.078; p<0.001), while no significant difference between Pitstop2 and DMSO was observed (Figure 2B). Cells treated with EIPA also showed little to no colocalization between tauPFFs and LAMP1, whereas treatment with DMSO and Pitstop2 demonstrated clear colocalization (Figure 2C). Significantly less tauPFFs colocalized with LAMP1 with EIPA treatment at 30 min (-6.413 ± 2.175; p=0.02), 1 hr (-14.65 ± 2.175; p=0.008) and 2 hrs (-21.53 ± 2.175; p<0.001) compared to DMSO vehicle control treatment (Figure 2D). No effect on colocalization between tauPFFs and LAMP1 was seen with Pitstop2 treatment (Figure 2D).

**Figure 2.**
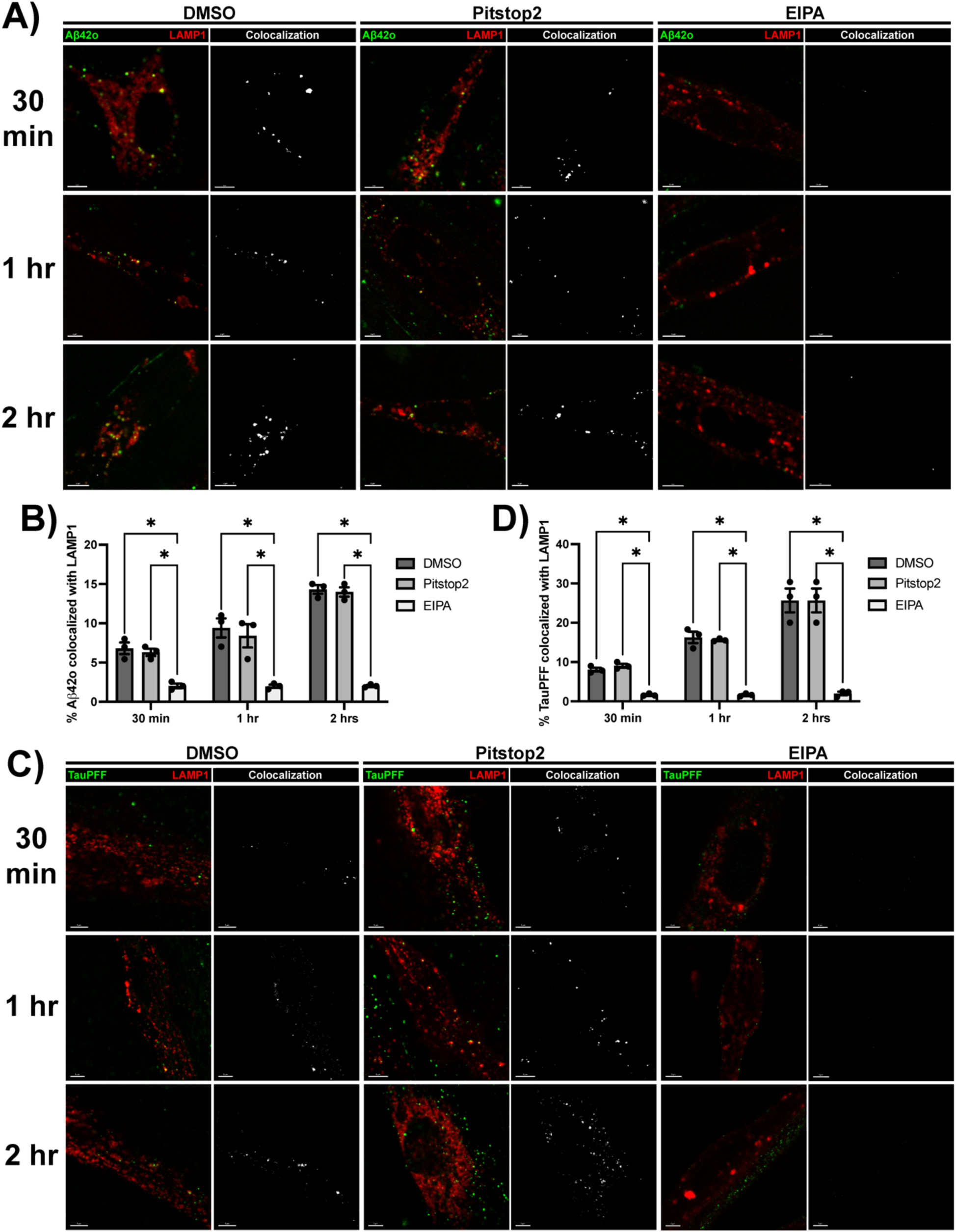
AB42o and tauPFF uptake is reduced with the inhibition of macropinocytosis. **A)** IPSC-derived neurons transduced with LAMP1-BFP (red) were treated with DMSO (vehicle control), Pitstop2 (CME inhibitor), or EIPA (macropinocytosis inhibitor). After treatment, neurons were incubated with Aβ42o (red) for 30 min, 1 hr, and 2 hrs. Colocalization between Aβ42o and LAMP1 was quantitated to measure uptake of oligomers to lysosomes (white) **B)** Quantification of the mean percent of Aβ42o signal colocalized with LAMP1 from three replicate experiments (n=3), each replicate containing at least 6 individual neuron cell body images at each time point for each treatment. Difference in colocalization between DMSO, Pitstop2 and EIPA was assessed using a two-way ANOVA with Sidak’s test. **C)** Neurons treated with DMSO, Pitstop2, or EIPA were incubated with tauPFFs (green) for 30 min, 1 hr, and 2 hrs. Colocalization between tauPFF and LAMP1-labeled lysosomes is marked by white pixels. **D)** Quantification of the mean percent of tauPFF colocalized with LAMP1 from three replicate experiments (n=3), each containing at least 6 individual neuron cell body images at each time point for each treatment. Difference between each treatment was assessed using a two-way ANOVA with Sidak’s test. *Data is presented as mean* ± *SEM. * p<0.05. Scale bar = 5μm*.

### Inhibition of Arf6, Rac1, Cdc42 and RhoA reduces the uptake of AB42o and tauPFFs to lysosomes

Having observed that EIPA treatment prevents the macropinocytosis of Aβ42 oligomers and tauPFFs to lysosomes, we next assessed the dependance of aggregate macropinocytosis on Arf6, Rac1, Cdc42 and RhoA activity. Neurons expressing LAMP1-BFP were treated with the small molecule inhibitors NAV-2729 (inhibits Arf6), EHT 1864 (inhibits Rac1), ML 141 (inhibits Cdc42), and Rhosin (inhibits RhoA). Following treatment, cells were incubated with tagged Aβ42o or tauPFFs for 30 minutes, after which the colocalization between each and LAMP1 was quantitated to measure uptake by macropinocytosis. In cells treated with DMSO vehicle control, both Aβ42o (Figure 3A) and tauPFFs (Figure 3C) were observed in LAMP-labeled lysosomes. However, treatment with all the inhibitors abolished uptake of both aggregates (Figure 3A&C). Measuring the percentage of Aβ42o colocalized with LAMP1 demonstrated significantly less colocalization compared to control after treatment with NAV-2729 (-7.456 ± 1.164; p<0.001), EHT 1864 (-7.820 ± 1.164; p<0.001), ML 141 (-7.913 ± 1.164; p<0.001) and Rhosin (-7.831 ± 1.164; p<0.001; Figure 3B). The same was demonstrated with the percentage of tauPFFs colocalized with LAMP1 after NAV-2729 (-11.86 ± 2.038; p=0.002), EHT 1864 (-11.50 ± 2.038; p=0.001), ML 141 (-11.88 ± 2.038; p=0.001) and Rhosin (-11.78 ± 2.038; p=0.001) treatment (Figure 3D).

**Figure 3.**
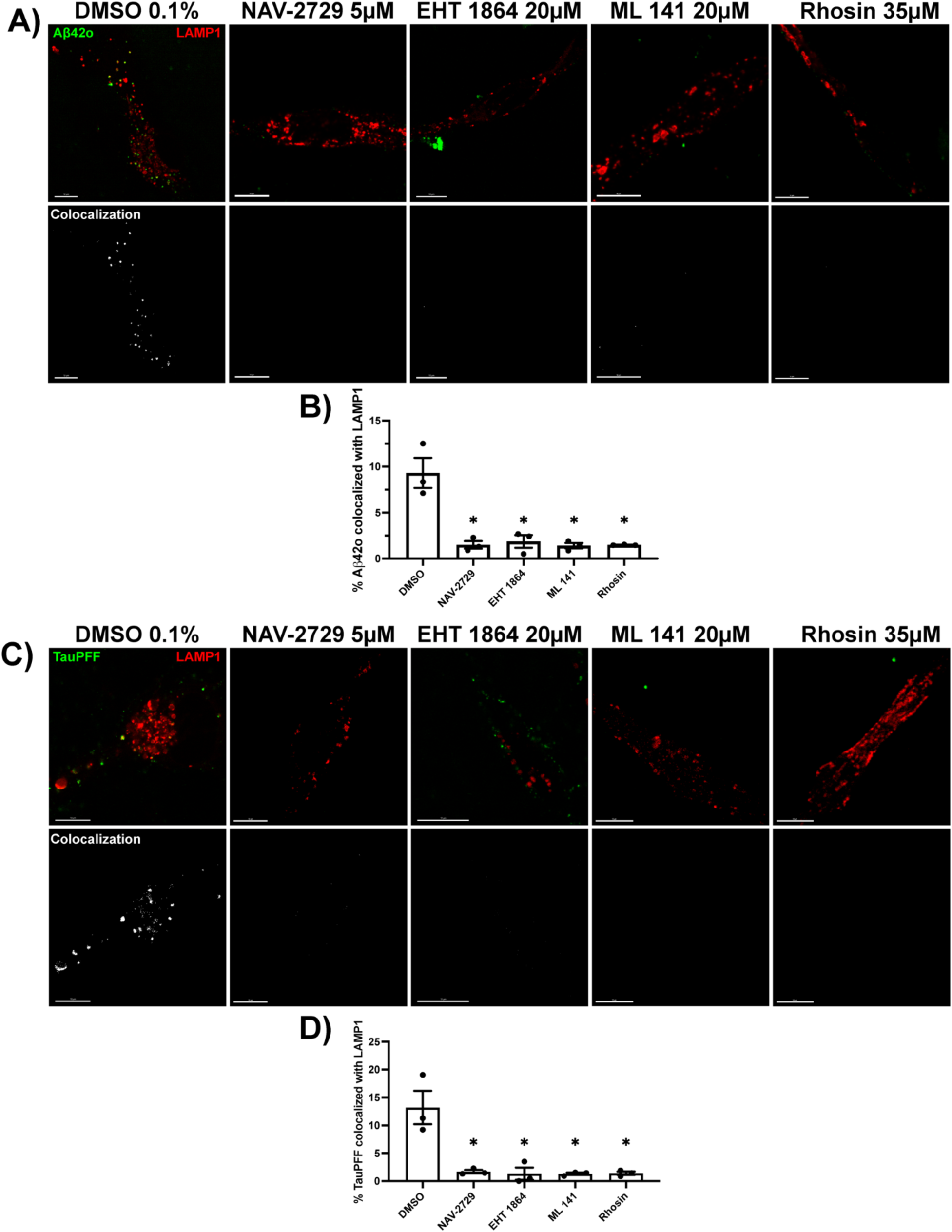
Small GTPase inhibition reduces the macropinocytosis of AB42o and tauPFFs to lysosomes. **A)** IPSC-derived neurons transduced with LAMP1-mCh (red) were treated with DMSO (vehicle control), NAV-2729 (Arf6 inhibitor), EHT 1864 (Rac1 inhibitor), ML 141 (Cdc42 inhibitor) or Rhosin (RhoA inhibitor). After treatment, neurons were incubated with Aβ42o (red) for 30 min the imaged live. Colocalization between Aβ42o and LAMP1 was quantitated with Imaris 10.1 (Bitplane) software to determine uptake of oligomers to lysosomes (white). **B)** Quantification of the mean percent of Aβ42o signal colocalized with LAMP1 from three replicate experiments (n=3), each containing at least 6 individual neuron cell body images at each time point for each treatment. Differences in colocalization between DMSO, NAV-2729, EHT 1864, ML 141, and Rhosin were assessed using a one-way ANOVA with Tukey’s test. **C)** Neurons treated with DMSO, NAV-2729, EHT 1864, ML 141, and Rhosin were incubated with tauPFFs (green) for 30 min, then imaged lived. Colocalization between tauPFFs and LAMP1-labeled lysosomes (white pixels) was measured. **D)** Quantification of the mean percent of tauPFF colocalized with LAMP1 from three replicate experiments (n=3), each containing at least 6 individual neuron cell body images at each time point for each treatment. Differences between each treatment were quantitated using a one-way ANOVA with Tukey’s test. *Data is presented as mean* ± *SEM. * p<0.05. Scale bar = 10μm*.

### Small GTPase inhibition reduces the co-uptake of AB42o or tau fibrils with dextran 70kDa

To further support that the inhibition of these GTPases prevents the uptake of Aβ42o by macropinocytosis, we examined the co-uptake of Aβ42o and the fluid phase marker dextran in the presence of these inhibitors. For these experiments, we utilized 70 kDa dextran (Dex70), as it has been demonstrated to be more selectively taken up by macropinocytosis than CME (39). Co-uptake was examined in neurons expressing LAMP1-BFP following treatment with the small GTPase inhibitors outlined above, as well as Pitstop2 and DMSO vehicle control. Treated cells were then incubated with Aβ42o and Dex70 for 30 minutes and then imaged. To measure changes in co-uptake, we first counted the number of Aβ42o and dextran puncta contained within the cell body of a neuron, then counted the number of puncta containing both Aβ42o and dextran (Aβ42o+/Dex70+). Bright Aβ42o and Dex70 puncta can be observed within cells treated with DMSO or Pitstop2 (Figure 4A). Cells treated with GTPase inhibitors show very few puncta within the cell body of neurons, and Aβ42o and Dex70 can be observed outlining some cell bodies (Figure 4A). Significantly lower numbers of Aβ42o puncta (p<0.001; Figure 4B), Dex70 puncta (p<0.001; Figure 4C), and Aβ42o+/Dex70+ puncta (p<0.001; Figure 4D) are observed within cells after treatment with NAV-2729, EHT 1864, ML 141, and Rhosin compared to either DMSO or Pitstop2 treatment. we also qualitatively examined whether Aβ42o is present in membrane ruffles containing actively polymerizing actin. To do so, neurons transduced with PLCδPH to mark membrane ruffles were incubated with oligomers for 10 mins, after which neurons were fixed and F-actin was labelled by fluorescently tagged phalloidin. After this incubation, Aβ42o can be observed within PLCδPH labeled F-actin containing protrusions from the membranes of cells (Figure 4E).

**Figure 4.**
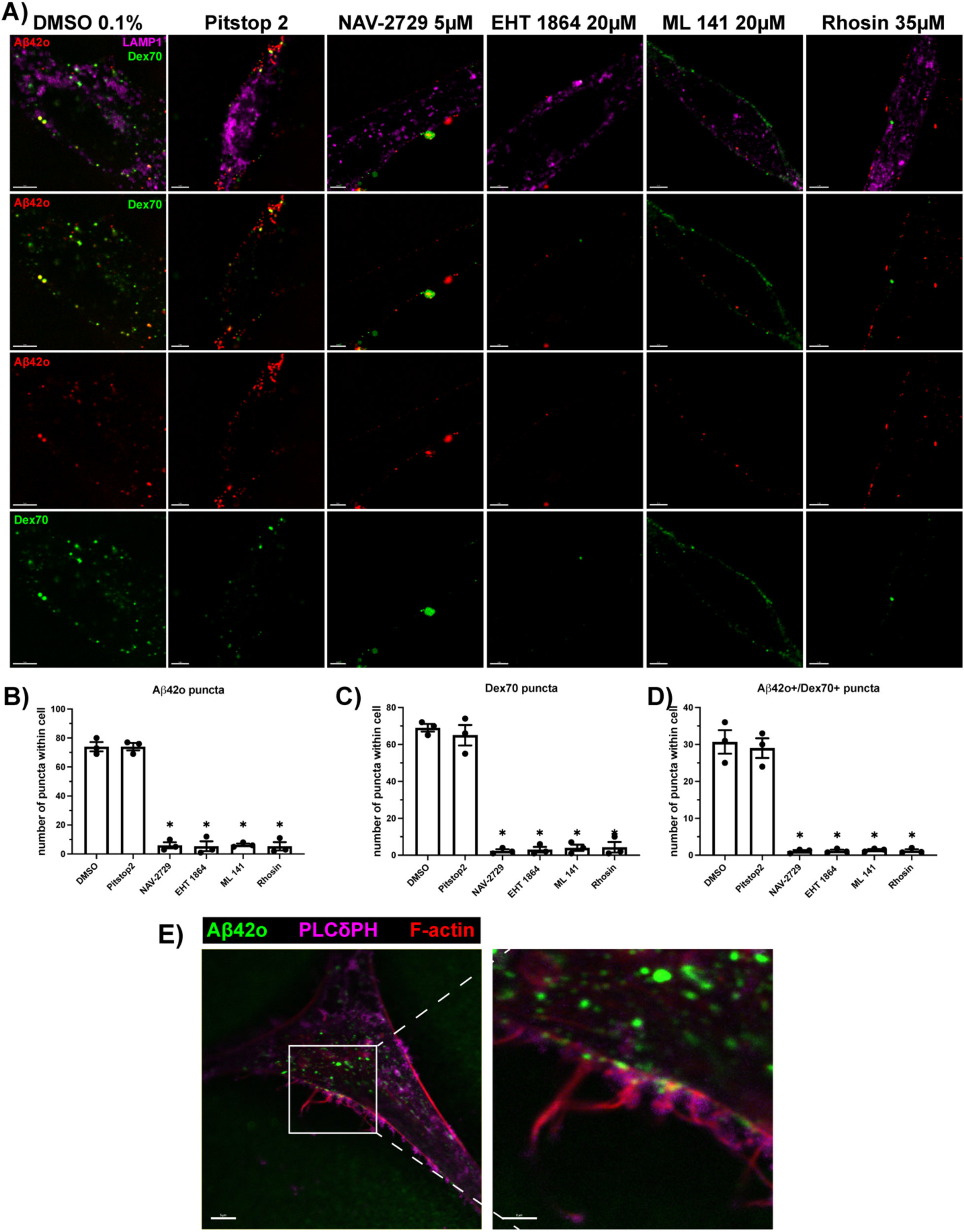
Small GTPase inhibition reduces co-uptake of AB42o and 70kDa dextran. **A)** IPSC-derived neurons transduced with LAMP1-mCh (magenta) were treated with DMSO (vehicle control), Pitstop2, NAV-2729, EHT 1864, ML 141 or Rhosin. After treatment, neurons were incubated with Aβ42o (red) and 70kDa dextran (Dex70; green) for 30 min then imaged live. To quantify the uptake of Aβ42o and Dex70, signal from each were used to create spots demarking puncta using Imaris. The number of puncta of each within the cell body of each neuron was counted. Puncta containing both Aβ42o and Dex70 (Aβ42o+/Dex70+) were also counted. Number of **B)** Aβ42o puncta, **C)** Dex70 puncta, and **D)** Aβ42o+/Dex70+ was quantified from three replicate experiments (n=3), each containing at least 6 individual neuron cell body images at each time point for each treatment. Differences in the number of puncta between DMSO, Pitstop2 NAV- 2729, EHT 1864, ML 141, and Rhosin were assessed using a one-way ANOVA with Tukey’s test. **D)** Image of a neuron transduced with PLC8PH (magenta) marking membrane ruffles and stained with fluorescently tagged phalloidin (red) marking F-actin. After 10 min incubation with Aβ42o (green), oligomers can be observed within membrane ruffles containing actively polymerizing actin. *Data are presented as mean ± SEM. * p<0.05. Scale bar = 5μm*.

This experiment was repeated with tauPFFs, and the co-uptake of tauPFFs with Dex70 was assessed. Similar to what was observed with Aβ42o, tauPFF and Dex70 are observed in neurons treated with DMSO or Pitstop2, while cells treated with small GTPase inhibitors showed little uptake of either tau PFFs or Dex70 (Figure 5A). The number of tauPFF puncta (Figure. 5B) or Dex70 puncta (Fibure 5C) within neurons was significantly lower with small GTPase inhibition (NAV-2729, EHT 1864, ML 141, or Rhosin) than with Pitstop2 or vehicle control (p<0.001). The same was observed when the number of puncta containing both tauPFFs and Dex70 were quantified (Figure 5D), with small GTPase inhibition significantly reducing the number of puncta within cells compared to CME inhibition (Pitstop2) or vehicle control (p<0.001). Like with Aβ42o, we qualitatively assessed tauPFFs in membrane ruffles containing actively polymerizing actin. Neurons transduced PLCδPH-BFP were incubated with tauPFFs for 10 mins, after which they were fixed, and F-actin was labelled by fluorescently tagged phalloidin. TauPFFs were observed within PLCδPH labeled membrane ruffles containing F-actin at the membranes of cells (Figure 5E).

**Figure 5.**
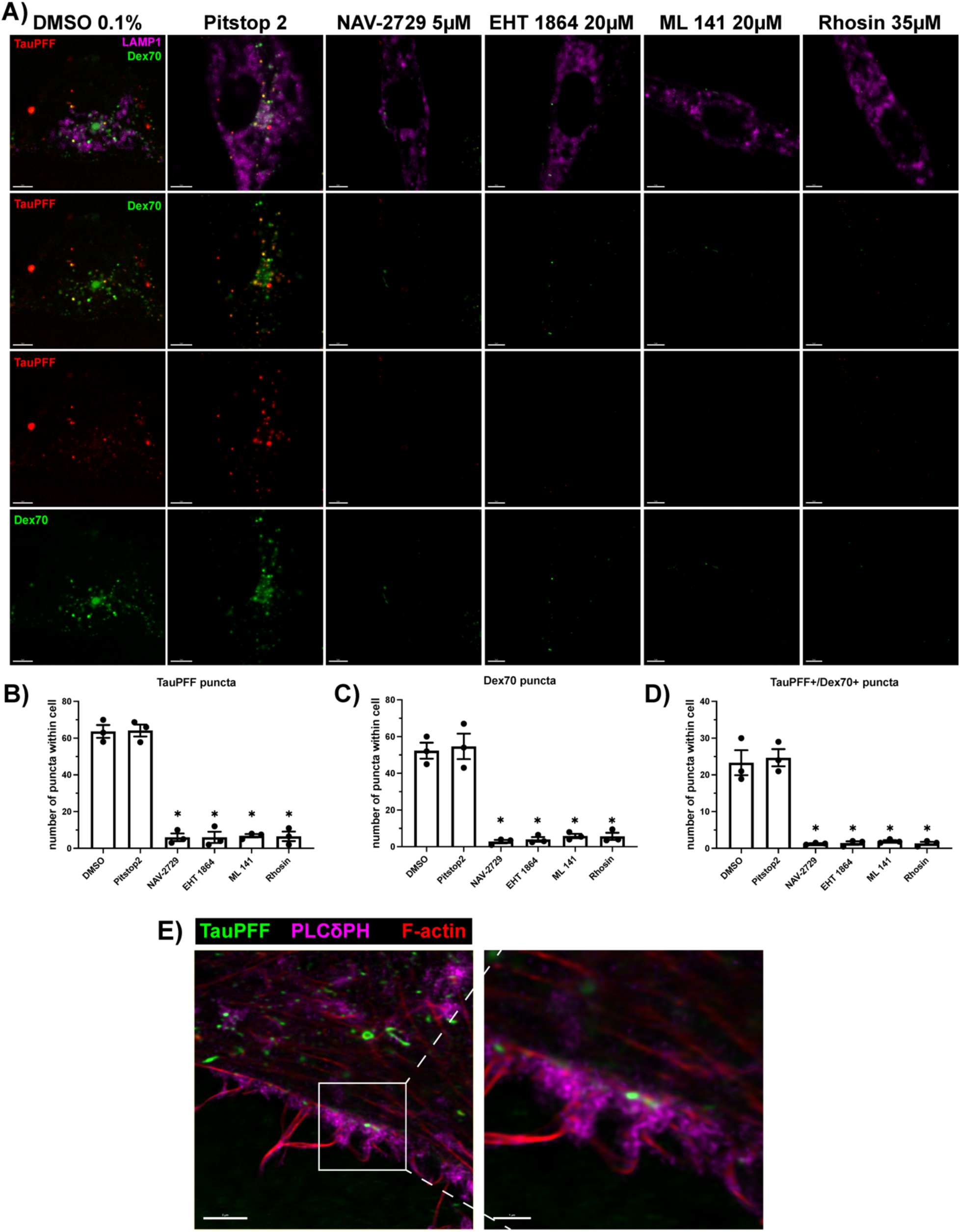
Small GTPase inhibition reduces co-uptake of tauPFFs and 70kDa dextran. **A)** IPSC-derived neurons transduced with LAMP1-mCh (magenta) were treated with DMSO (vehicle control), Pitstop2, NAV-2729, EHT 1864, ML 141 or Rhosin. After treatment, neurons were incubated with tauPFFs (red) and 70kDa dextran (Dex70; green) for 30 min the imaged live. To quantify the uptake of tauPFFs and Dex70, signal from each were used to create spots demarking puncta using Imaris. Puncta containing both tauPFFs and Dex70 (TauPFF+/Dex70+) were also counted. The number of **B)** TauPFF puncta, **C)** Dex70 puncta, and **D)** TauPFF+/Dex70+ puncta within neuron cell bodies were quantified from from three replicate experiments (n=3), each containing at least 6 individual neuron cell body images at each time point for each treatment. Differences in the number of puncta between DMSO, Pitstop2 NAV-2729, EHT 1864, ML 141, and Rhosin were assessed using a one-way ANOVA with Tukey’s test. **E)** Image of a neuron transduced with PLC8PH (magenta) marking membrane ruffles and stained with fluorescently tagged phalloidin (red) marking F-actin. After 10 min incubation with tauPFFs (green), they can be observed within membrane ruffles containing actively polymerizing actin. *Data are presented as mean ± SEM. * p<0.05. Scale bar = 5μm*.

### AB42 oligomers and tau fibrils increase the amount of active Rac1, Cdc42, and RhoA

Next, we examined if the incubation of Aβ42o or tauPFFs with cells increases the amount of active GTP-bound Rac1, Cdc42, and RhoA. To do so, we incubated N2a cells with no treatment (untreated), Aβ42o, tauPFFs, or EGF as a positive control. Cells were then lysed, and a Rac1, Cdc42, or RhoA G-LISA assays was used to quantify the amount of GTP-bound protein within the lysate. In response to Aβ42o, tauPFF or EGF treatment, levels of GTP-bound Rac1 were significantly higher than in untreated N2a cells (p<0.001; Figure 6A). The amount of active GTP-bound Cdc42 was also elevated in response to Aβ42o, tauPFFs or EGF treatment (p<0.001; Figure 6B). Finally, increased activity of RhoA was demonstrated by increase GTP-bound RhoA in N2a cells incubated with Aβ42o, tauPFFs and EGF (p<0.001; Figure 6C).

**Figure 6.**
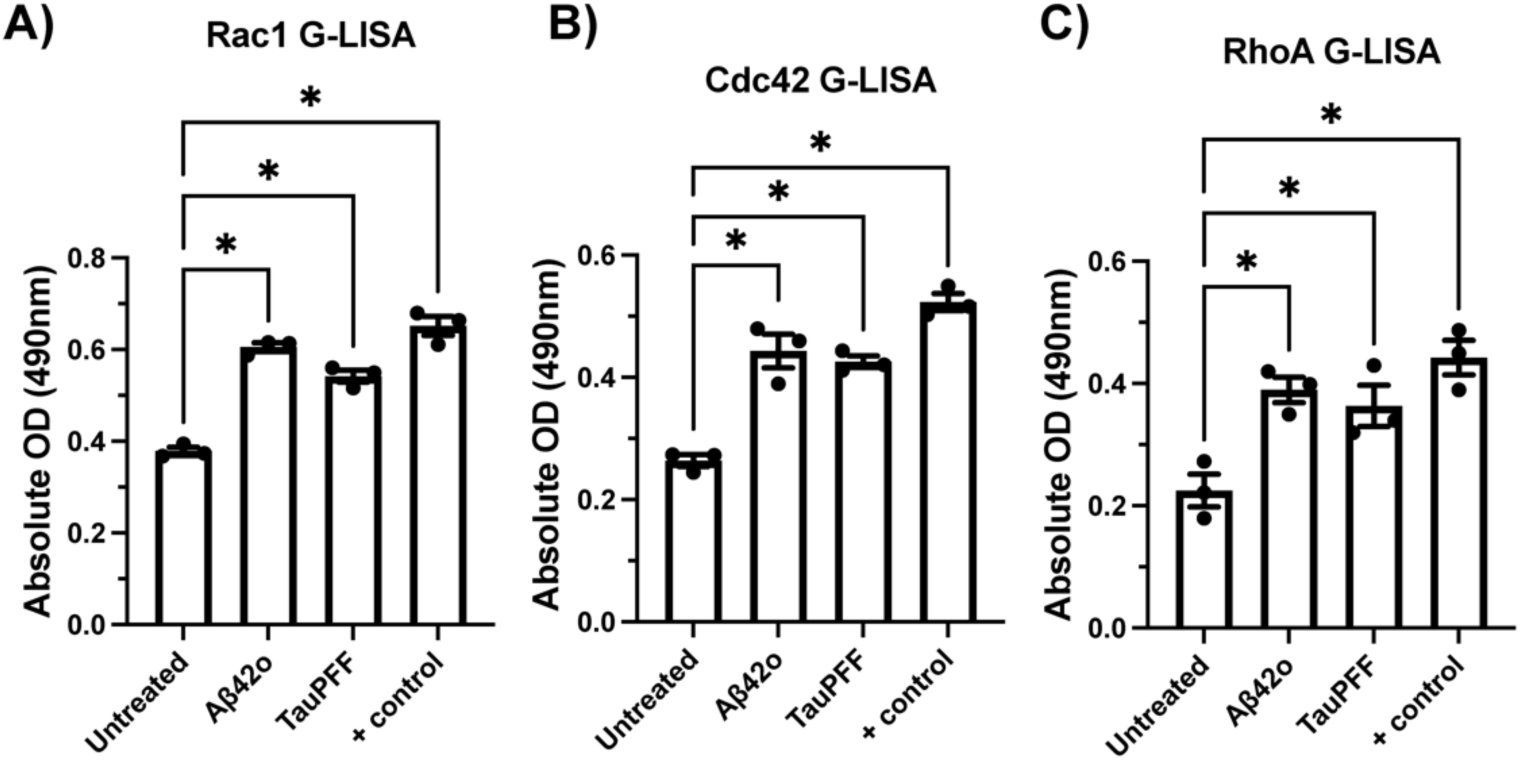
**AB42o or tauPFF incubation increases Rac1, Cdc42 and RhoA activity in Neuro2a cells.** Neuro2a (N2a) mouse neuroblastoma cells were incubated with Aβ42o, tauPFFs or EGF (positive control) while on ice, then incubated for 5 minutes at 37°C and 5% CO_2_ to allow for the initiation of macropinocytosis. Cells were moved back to ice following this incubation and lysates were immediately collected and used for Rac1, Cdc42 or RhoA G-LISA activation assays. To measure the relative amount of active GTP-bound levels of each protein, the absolute optical density (OD) was measured in lysate loaded into G-LISA assay wells excited with a 490nm laser. Absolute OD was measured in lysates from untreated cells, or cells treated with Aβ42o, tauPFFs, and EGF and levels of **A)** GTP-bound Rac1, **B)** GTP-bound Cdc42, and **C)** GTP-bound RhoA were compared. Each of these experiments was ran in technical triplicates, and three biological replicates were performed (n=3). The difference in absolute OD was compared to untreated control by a one-way ANOVA with Tukey’s test. *Data are presented as mean ± SEM. * p<0.05*.

### Uptake of aggregates to lysosomes results in the accumulation of intracellular tau in lysosomes

Finally, we sought to characterize the consequences of Aβ42o and tauPFF uptake by human neurons on tau localization. To investigate this, neurons were transduced with LAMP1- BFP as well as tau containing a Halo-tag sequence (Halo-tau). Cells were then incubated with no aggregates, Aβ42o alone, tau PFFs alone, or both Aβ42o and tauPFFs together. Following incubation, Halo-tau was tagged with a fluorescent Halo-tag ligand, labeling all of the Halo-tau present in cells at that given time. Neurons were then imaged live immediately following Halo-tag labelling (0 hours) or imaged 6 hours, 24 hours, 48 hours, 5 days, or 7 days following Halo-tag labeling. Halo-tau appeared entirely cytosolic at 0 hours and 6 hours, with small amounts colocalized with lysosomes (Figure 7A). However, by 24 and 48 hours, in lysosomes containing Aβ42o and tauPFFs, accumulation of Halo-tau can be observed but remains mostly cytoplasmic in the absence of aggregates (Figure 7A). By 5 and 7 days, further accumulation of Halo-tau is observed in cells containing Aβ42o and tauPFFs, with less remaining in the cytoplasm compared to untreated (Figure 7A). Quantifying the percent of labeled Halo-tau colocalized with LAMP1, the first significant increase in colocalization is observed at 24 hours in cells loaded with both aggregates (p<0.05; Figure 7B), with the difference between all aggregate loaded conditions and untreated cells increasing at 48 hours, 5 days, and 7 days (p<0.05; Figure 7B). At 5 and 7 days, the mean percent colocalized is highest in cells loaded with both Aβ42o and tauPFFs, although it is not significantly higher than with Aβ42o or tauPFFs alone (Figure 7B).

**Figure 7.**
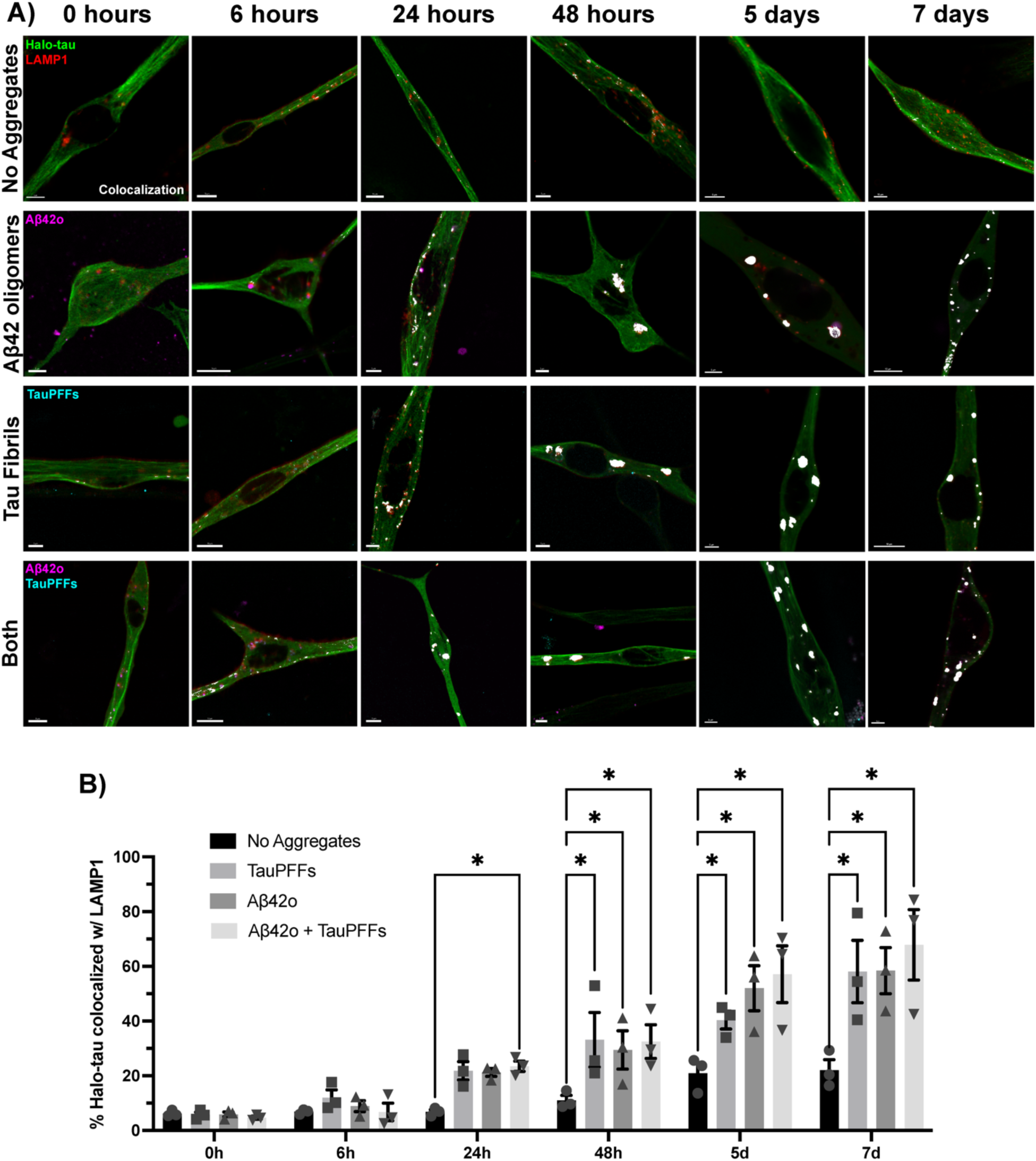
Uptake of aggregates to lysosomes results in the accumulation of cytoplasmic tau within lysosomes. **A)** IPSC-derived neurons were transduced with Halo-tagged human tau (Halo-tau) and LAMP1- BFP (red). Transduced neurons were then incubated with no aggregates, Aβ42o alone (magenta), tauPFFs alone (cyan), or both Aβ42o and tauPFFs together for 18 hours (overnight). Excess aggregates were washed off following loading, and Halo-tau was labeled using JaneliaFluor- labeled Halo-tag ligand (green). Neurons were then imaged live directly after this labeling (0 hrs), or imaged at 6 hours, 24 hours, 48 hours, 5 days, and 7 days after Halo-tag labeling. Colocalization between labeled Halo-tau and LAMP1-mCh was quantitated (white pixels). **B)** The mean percentage of Halo-tau colocalized with LAMP1 was analyzed from three replicate experiments (n=3), each containing at least 6 individual neuron cell body images at each time point for each treatment. Significant differences between each condition within a time point were analyzed using a two-way ANOVA with Sidak’s test. *Data are presented as mean ± SEM. * p<0.05. Scale bar = 5um*.

## Discussion

The objective of the current study was to build upon previous observations of the uptake of Aβ and tau aggregates by macropinocytosis and examine if this mechanism requires the activity of small GTPases in cultured human neurons. We first demonstrated that the uptake of Aβ and tau aggregates to lysosomes is reduced by treatment with the macropinocytosis inhibitor EIPA but not inhibited by the CME inhibitor Pitstop2. Then, we demonstrated that the macropinocytosis of Aβ42 oligomers and tau fibrils requires the activity of Arf6, Rac1, Cdc42 and RhoA by inhibition with small molecule inhibitors. To further inhibition resulted in reduced uptake through preventing macropinocytosis, we demonstrated that treatment with inhibitors reduced the co-uptake of oligomers and fibrils with a macropinocytosis selective fluid phase marker 70kDa dextran (39). Additionally, both Aβ42 oligomers and tau fibrils were found in membrane ruffles containing F-actin on the membrane of human neurons. To demonstrate that these observations were directly the result of incubating cells with aggregates increases in Rac1, Cdc42, and RhoA activity were observed in response to incubation with N2a cells. Finally, to characterize the physiological consequence of Aβ42 oligomer and tau fibrils uptake, we demonstrated the abnormal accumulation of cytoplasmic tau in lysosomes containing extracellular aggregates similar to observations from a previous study (40).

Investigating the mechanisms that mediate the transcellular spread of protein aggregates in AD has begun to attract significant attention for its like role in the propagation of Aβ and tau pathology leading to progression of the disease. Prion-like mechanisms in AD have been proposed, by which aggregated Aβ or tau released by one neuron can be taken up by an adjacent or connected neuron (13). Internalized aggregates can then mediate aggregation in this new neuron through direct nucleation of aggregation or inducing dysfunction in the cells ability to degrade aggregated protein (40). Subsequent release of aggregates further propagates regional aggregation, which is thought to underly the spreading of pathology between structurally connected brain regions (13). Thus, elucidating the specific mechanisms by which one neuron can take up Aβ or tau aggregates could provide new therapeutic targets aimed at stopping the propagation of Aβ and tau aggregation in AD. Macropinocytosis was suggested as a mechanism of interest in this model, as its ability to take up large amount of extracellular fluid including its contents would be suited for the uptake of soluble Aβ and tau aggregates (24,41). So far, macropinocytosis has been demonstrated to play a role in both Aβ and tau aggregate uptake in-vitro using compounds which block aspects of macropinocytosis. This has been demonstrated by inhibition of actin polymerization (42), which macropinocytosis is dependent on. The PI3K inhibitor wortmannin has also been used to prevent aggregate uptake (26), which prevents the necessary conversion of PI(4,5)P_2_ to PIP_3_ required for macropinocytic cup closure (43). Studies have also demonstrated reduced uptake in cells treated with amiloride, a commonly used inhibitor of macropinocytosis (26). Amiloride and its derivatives, like EIPA, have been shown to mediate this effect on macropinocytosis by blocking Na+/H+ exchangers at the plasma membrane, and as a result reduce the local activity of Rac1 and Cdc42 through pH disruptions (32). Like these previous studies, we observed the same effect of amiloride treatment on aggregate uptake and demonstrated the same effect occurs with the inhibition of the small GTPases Arf6, Rac1, Cdc42 and RhoA directly. The evidence provided in the current study further supports these previous observations and highlights the requirement of the activity of these GTPases in the uptake of Aβ and tau aggregates.

Previous studies have compared the endocytosis mechanisms utilized by monomeric and aggregated Aβ and tau species. Extracellular monomeric Aβ42 has shown an increased preference for internalization by macropinocytosis compared to Aβ40 (44). It was also found to endocytose at a significantly higher rate (44). Differences in tau species uptake have also been investigated but have yielded conflicting findings. In one study, human neurons took up aggregated tau primarily through CME, and monomeric tau through macropinocytosis (42). Supporting this, aggregated tau uptake was shown to require dynamin activity, and inhibition of dynamin with the inhibitor dynasore prevented tau aggregate uptake by clathrin-dependent endocytosis (45). The use of dynasore to selectively inhibit CME is worth reconsidering, given it has been shown to result in off-target effects on membrane cholesterol levels and can inhibit macropinocytosis (34). However, other studies have demonstrated that tau aggregates such as fibrils are taken up by macropinocytosis and requires HSPGs (27,46). The results of this study should be considered specific to the uptake of recombinant Aβ42 oligomers and tau P301S mutant fibrils, and future work should characterize the exact species and size of aggregates internalized by macropinocytosis and demonstrate its regulation by the small GTPases examined here. Exploring the uptake of patient-derived or AD mouse model-derived Aβ and tau aggregates is another consideration worth examining in future studies.

With many attempts at targeting the production of Aβ or at clearing Aβ and tau aggregates to treat AD yielding minimal progress towards meaningful new therapeutic options for patients, there is an increasing push to identify unexplored disease mechanisms. Preventing the propagation of Aβ and tau aggregation has been proposed as a mechanism that could be targeted to halt the progression of AD (14). Here, we show that inhibitors of Arf6, Rac1, Cdc42 and RhoA control the uptake of Aβ42 oligomers and tau fibrils. Modulating their activity could hold significant therapeutic promise, given there is evidence that their activity might be dysregulated in AD. Increased activity of the Rac1 effector p21-activated kinase 1 (PAK1) at the plasma membrane, as well as increased membrane localization of Rac1, has been observed in both AD patient brain samples and AD mouse models (47). Increased RhoA activity has been observed in the neurites of degenerating neurons in APP-overexpressing AD mouse models (48). Alterations in endocytosis have been observed in both AD and normal aging have been identified, with increases in APP endocytosis and its cleavage to form Aβ implicated in both (49,50). Future studies which contribute to characterizing how macropinocytosis and other endocytic mechanisms contribute to both the production and propagation of Aβ and tau aggregates may allow for the therapeutic targeting of one or more of these mechanisms and halt the progression of AD.

## Conclusion

Previous work has implicated macropinocytosis in the uptake of extracellular AB and tau aggregates. However, the exact regulatory proteins involved have previously not been identified. Here, we provide evidence for the requirement of Arf6, Rac1, Cdc42 and RhoA activity in the macropinocytosis of Aβ42 oligomers and tau PFFs and their delivery to lysosomes. Further, the uptake of these aggregates to lysosomes resulted in the abnormal accumulation of cytoplasmic tau within aggregate-containing lysosomes. Demonstrating this provides supports for previous observations of the macropinocytosis of AD aggregates, and its potential role in transcellular spread of protein aggregation. Targeting the activity of these GTPases may hold promise as a novel AD therapy by preventing the uptake of aggregates in neurons, halting the spread of Aβ and tau aggregates from neuron-to-neuron.

## References

1. Braak H, Braak E. Neuropathological stageing of Alzheimer-related changes. Acta Neuropathol. 1991 Sep;82(4):239–59.

2. Alzheimer’s Association. 2016 Alzheimer’s disease facts and figures. Alzheimer’s & Dementia. 2016 Apr;12(4):459–509.

3. Kelley BJ, Petersen RC. Alzheimer’s Disease and Mild Cognitive Impairment. Neurologic Clinics. 2007 Aug;25(3):577–609.

4. Blennow K, de Leon MJ, Zetterberg H. Alzheimer’s disease. The Lancet. 2006 Jul;368(9533):387–403.

5. Villemagne VL, Burnham S, Bourgeat P, Brown B, Ellis KA, Salvado O, et al. Amyloid β deposition, neurodegeneration, and cognitive decline in sporadic Alzheimer’s disease: a prospective cohort study. The Lancet Neurology. 2013 Apr;12(4):357–67.

6. Bateman RJ, Xiong C, Benzinger TLS, Fagan AM, Goate A, Fox NC, et al. Clinical and Biomarker Changes in Dominantly Inherited Alzheimer’s Disease. N Engl J Med. 2012 Aug 30;367(9):795–804.

7. Bennett RE, DeVos SL, Dujardin S, Corjuc B, Gor R, Gonzalez J, et al. Enhanced Tau Aggregation in the Presence of Amyloid β. The American Journal of Pathology. 2017 Jul;187(7):1601–12.

8. Walsh DM, Selkoe DJ. Deciphering the Molecular Basis of Memory Failure in Alzheimer’s Disease. Neuron. 2004 Sep;44(1):181–93.

9. Gouras GK, Tampellini D, Takahashi RH, Capetillo-Zarate E. Intraneuronal β-amyloid accumulation and synapse pathology in Alzheimer’s disease. Acta Neuropathol. 2010 May;119(5):523–41.

10. Ittner LM, Götz J. Amyloid-β and tau — a toxic pas de deux in Alzheimer’s disease. Nat Rev Neurosci. 2011 Feb;12(2):67–72.

11. Ballatore C, Lee VMY, Trojanowski JQ. Tau-mediated neurodegeneration in Alzheimer’s disease and related disorders. Nat Rev Neurosci. 2007 Sep;8(9):663–72.

12. Masters CL, Bateman R, Blennow K, Rowe CC, Sperling RA, Cummings JL. Alzheimer’s disease. Nat Rev Dis Primers. 2015 Oct 15;1(1):15056.

13. Soto C, Pritzkow S. Protein misfolding, aggregation, and conformational strains in neurodegenerative diseases. Nat Neurosci. 2018 Oct;21(10):1332–40.

14. Vaquer-Alicea J, Diamond MI. Propagation of Protein Aggregation in Neurodegenerative Diseases. Annu Rev Biochem. 2019 Jun 20;88(1):785–810.

15. Kerr MC, Teasdale RD. Defining Macropinocytosis. Traffic. 2009 Apr;10(4):364–71.

16. Buckley CM, King JS. Drinking problems: mechanisms of macropinosome formation and maturation. FEBS J. 2017 Nov;284(22):3778–90.

17. Donaldson JG. Macropinosome formation, maturation and membrane recycling: lessons from clathrin-independent endosomal membrane systems. Phil Trans R Soc B. 2019 Feb 4;374(1765):20180148.

18. Burton KM, Johnson KM, Krueger EW, Razidlo GL, McNiven MA. Distinct forms of the actin cross-linking protein α-actinin support macropinosome internalization and trafficking. Barber D, editor. MBoC. 2021 Jul 15;32(15):1393–407.

19. Palm W. Metabolic functions of macropinocytosis. Phil Trans R Soc B. 2019 Feb 4;374(1765):20180285.

20. Donaldson J, Poratshliom N, Cohen L. Clathrin-independent endocytosis: A unique platform for cell signaling and PM remodeling. Cellular Signalling. 2009 Jan;21(1):1–6.

21. Jin J, Shen Y, Zhang B, Deng R, Huang D, Lu T, et al. In situ exploration of characteristics of macropinocytosis and size range of internalized substances in cells by 3D-structured illumination microscopy. IJN. 2018 Sep;Volume 13:5321–33.

22. Lim JP, Gleeson PA. Macropinocytosis: an endocytic pathway for internalising large gulps. Immunol Cell Biol. 2011 Nov;89(8):836–43.

23. Bloomfield G, Kay RR. Uses and abuses of macropinocytosis. Journal of Cell Science. 2016 Jan 1;jcs.176149.

24. Zeineddine R, Yerbury JJ. The role of macropinocytosis in the propagation of protein aggregation associated with neurodegenerative diseases. Front Physiol [Internet]. 2015 Oct 16 [cited 2023 Jan 27];6. Available from: http://journal.frontiersin.org/Article/10.3389/fphys.2015.00277/abstract

25. Yerbury JJ. Protein aggregates stimulate macropinocytosis facilitating their propagation. Prion. 2016 Mar 3;10(2):119–26.

26. Nazere K, Takahashi T, Hara N, Muguruma K, Nakamori M, Yamazaki Y, et al. Amyloid Beta Is Internalized via Macropinocytosis, an HSPG- and Lipid Raft-Dependent and Rac1- Mediated Process. Front Mol Neurosci. 2022 Feb 11;15:804702.

27. Holmes BB, DeVos SL, Kfoury N, Li M, Jacks R, Yanamandra K, et al. Heparan sulfate proteoglycans mediate internalization and propagation of specific proteopathic seeds. Proc Natl Acad Sci USA [Internet]. 2013 Aug 13 [cited 2023 Jan 27];110(33). Available from: https://pnas.org/doi/full/10.1073/pnas.1301440110

28. Egami Y, Taguchi T, Maekawa M, Arai H, Araki N. Small GTPases and phosphoinositides in the regulatory mechanisms of macropinosome formation and maturation. Front Physiol [Internet]. 2014 Sep 30 [cited 2023 Jan 27];5. Available from: http://journal.frontiersin.org/article/10.3389/fphys.2014.00374/abstract

29. Buee L, Bussiere T, Buee-Scherrer V, Delacourte A, Hof PR. Tau protein isoforms, phosphorylation and role in neurodegenerative disorders. Brain Research Reviews. 2000;

30. Dahlgren KN, Manelli AM, Stine WB, Baker LK, Krafft GA, LaDu MJ. Oligomeric and Fibrillar Species of Amyloid-β Peptides Differentially Affect Neuronal Viability*. Journal of Biological Chemistry. 2002 Aug 30;277(35):32046–53.

31. Stine WB, Dahlgren KN, Krafft GA, LaDu MJ. *In Vitro* Characterization of Conditions for Amyloid-β Peptide Oligomerization and Fibrillogenesis*. Journal of Biological Chemistry. 2003 Mar 28;278(13):11612–22.

32. Koivusalo M, Welch C, Hayashi H, Scott CC, Kim M, Alexander T, et al. Amiloride inhibits macropinocytosis by lowering submembranous pH and preventing Rac1 and Cdc42 signaling. Journal of Cell Biology. 2010 Feb 22;188(4):547–63.

33. von Kleist L, Stahlschmidt W, Bulut H, Gromova K, Puchkov D, Robertson MJ, et al. Role of the clathrin terminal domain in regulating coated pit dynamics revealed by small molecule inhibition. Cell. 2011 Aug 5;146(3):471–84.

34. Preta G, Cronin JG, Sheldon IM. Dynasore - not just a dynamin inhibitor. Cell Commun Signal. 2015 Dec;13(1):24.

35. Chiu J, Krupa JM, Seah C, Pasternak SH. Small GTPases control macropinocytosis of amyloid precursor protein and cleavage to amyloid-β. Heliyon. 2024 May;10(10):e31077.

36. Yu Q, Gratzke C, Wang R, Li B, Kuppermann P, Herlemann A, et al. A NAV2729-sensitive mechanism promotes adrenergic smooth muscle contraction and growth of stromal cells in the human prostate. Journal of Biological Chemistry. 2019 Aug;294(32):12231–49.

37. Francis TC, Gaynor A, Chandra R, Fox ME, Lobo MK. The Selective RhoA Inhibitor Rhosin Promotes Stress Resiliency Through Enhancing D1-Medium Spiny Neuron Plasticity and Reducing Hyperexcitability. Biological Psychiatry. 2019 Jun;85(12):1001–10.

38. Lorenzen A, Samosh J, Vandewark K, Anborgh PH, Seah C, Magalhaes AC, et al. Rapid and Direct Transport of Cell Surface APP to the Lysosome defines a novel selective pathway. Molecular Brain. 2010;3(11).

39. Li L, Wan T, Wan M, Liu B, Cheng R, Zhang R. The effect of the size of fluorescent dextran on its endocytic pathway: Size-based endocytic entry for fluid cargoes. Cell Biol Int. 2015 May;39(5):531–9.

40. Piovesana E, Magrin C, Ciccaldo M, Sola M, Bellotto M, Molinari M, et al. Tau accumulation in degradative organelles is associated to lysosomal stress. Sci Rep. 2023 Oct 21;13(1):18024.

41. Jones AT. Macropinocytosis: searching for an endocytic identity and role in the uptake of cell penetrating peptides. J Cellular Mol Med. 2007 Jul;11(4):670–84.

42. Evans LD, Wassmer T, Fraser G, Smith J, Perkinton M, Billinton A, et al. Extracellular Monomeric and Aggregated Tau Efficiently Enter Human Neurons through Overlapping but Distinct Pathways. Cell Reports. 2018 Mar;22(13):3612–24.

43. Yoshida S, Hoppe AD, Araki N, Swanson JA. Sequential signaling in plasma-membrane domains during macropinosome formation in macrophages. Journal of Cell Science. 2009 Sep 15;122(18):3250–61.

44. Wesén E, Jeffries GDM, Matson Dzebo M, Esbjörner EK. Endocytic uptake of monomeric amyloid-β peptides is clathrin- and dynamin-independent and results in selective accumulation of Aβ(1–42) compared to Aβ(1–40). Sci Rep. 2017 May 17;7(1):2021.

45. Wu JW, Herman M, Liu L, Simoes S, Acker CM, Figueroa H, et al. Small Misfolded Tau Species Are Internalized via Bulk Endocytosis and Anterogradely and Retrogradely Transported in Neurons. Journal of Biological Chemistry. 2013 Jan;288(3):1856–70.

46. Stopschinski BE, Holmes BB, Miller GM, Manon VA, Vaquer-Alicea J, Prueitt WL, et al. Specific glycosaminoglycan chain length and sulfation patterns are required for cell uptake of tau versus α-synuclein and β-amyloid aggregates. Journal of Biological Chemistry. 2018 Jul;293(27):10826–40.

47. Ma QL, Yang F, Calon F, Ubeda OJ, Hansen JE, Weisbart RH, et al. p21-activated Kinase- aberrant Activation and Translocation in Alzheimer Disease Pathogenesis. Journal of Biological Chemistry. 2008 May;283(20):14132–43.

48. Huesa G, Baltrons MA, Gómez-Ramos P, Morán A, García A, Hidalgo J, et al. Altered Distribution of RhoA in Alzheimer’s Disease and AβPP Overexpressing Mice. Lovell MA, editor. JAD. 2010 Jan 6;19(1):37–56.

49. Burrinha T, Guimas Almeida C. Aging impact on amyloid precursor protein neuronal trafficking. Current Opinion in Neurobiology. 2022 Apr;73:102524.

50. Burrinha T, Martinsson I, Gomes R, Terrasso AP, Gouras GK, Almeida CG. Upregulation of APP endocytosis by neuronal aging drives amyloid-dependent synapse loss. Journal of Cell Science. 2021 May 1;134(9):jcs255752.

